# Perception of a conserved family of plant signalling peptides by the receptor kinase HSL3

**DOI:** 10.1101/2021.10.25.465685

**Authors:** Jack Rhodes, Andra-Octavia Roman, Marta Bjornson, Benjamin Brandt, Paul Derbyshire, Michele Wyler, Marc Schmid, Frank L.H. Menke, Julia Santiago, Cyril Zipfel

## Abstract

Plant genomes encode hundreds of secreted peptides; however, relatively few have been characterised. We report here an uncharacterised, stress-induced family of plant signalling peptides, which we call CTNIPs. Based on the role of the common co-receptor BRASSINOSTEROID INSENSITIVE 1-ASSOCIATED KINASE 1 (BAK1) in CTNIP-induced responses, we identified the orphan receptor kinase HAESA-LIKE 3 (HSL3) as the CTNIP receptor via a proteomics approach. CTNIP binding, ligand-triggered complex formation with BAK1, and induced downstream responses all involve HSL3. Notably, the HSL3-CTNIP signalling module is evolutionarily ancient, predating the divergence of extant angiosperms. The identification of this signalling module will help establish its physiological role and provides a resource to understand further receptor-ligand co-evolution.

## Introduction

Secreted plant peptides play major roles in growth, development and stress responses (Olsson et al., 2019). Whilst many hundreds of peptides are predicted to be encoded in plant genomes, relatively few have been characterised and their corresponding receptors are mostly unknown (Olsson et al., 2019).

Most signalling peptides are recognised by cell-surface localised receptors, especially by leucine-rich repeat receptor kinases (LRR-RKs). LRR-RKs generally function through the ligand-dependent recruitment of a shape complementary co-receptor to form an active signalling complex (Hohmann et al., 2017). The best characterised peptide receptors belong to LRR-RK subfamily XI, which recognize distinct families of plant peptides involved in growth, development or stress responses (Furumizu et al., 2021). Notably, the LRR-RK MIK2, which belongs to the closely related LRR-RK subfamily XIIb (an outgroup recently included within subfamily XI; (Liu et al., 2017; Man et al., 2020)) was recently shown to perceive SCOOP peptides (Hou et al., 2021; Rhodes et al., 2021). Despite intensive studies on the LRR-RK subfamily XI, the ligand for HAESA-like 3 (HSL3) has remained elusive, hindering our ability to investigate peptide-receptor coevolution across the family (Furumizu et al., 2021; Lee et al., 2020).

## Results and discussion

Several peptides (PEPs, PIPs, SCOOPs, CLEs and IDLs) recognised by LRR-RKs from subfamily XI or XIIb are transcriptionally up-regulated by abiotic or biotic stresses (Bartels et al., 2013; Gully et al., 2019; Kim et al., 2021; Takahashi et al., 2018; Vie et al., 2015). In order to identify novel stress-induced signalling peptides, we searched for *Arabidopsis thaliana* (hereafter, Arabidopsis) transcripts encoding short proteins (<150 amino acids) with a predicted signal peptide, which were induced upon biotic elicitor treatment (Bjornson et al., 2021). Through this analysis, we identified an uncharacterised family of peptides with 5 predicted members, which we named CTNIP1 to 5 (pronounced catnip) based on relatively conserved residues within the peptides (Figure 1a-b; Figure 1-figure supplement 1a-b).

**Figure 1.**
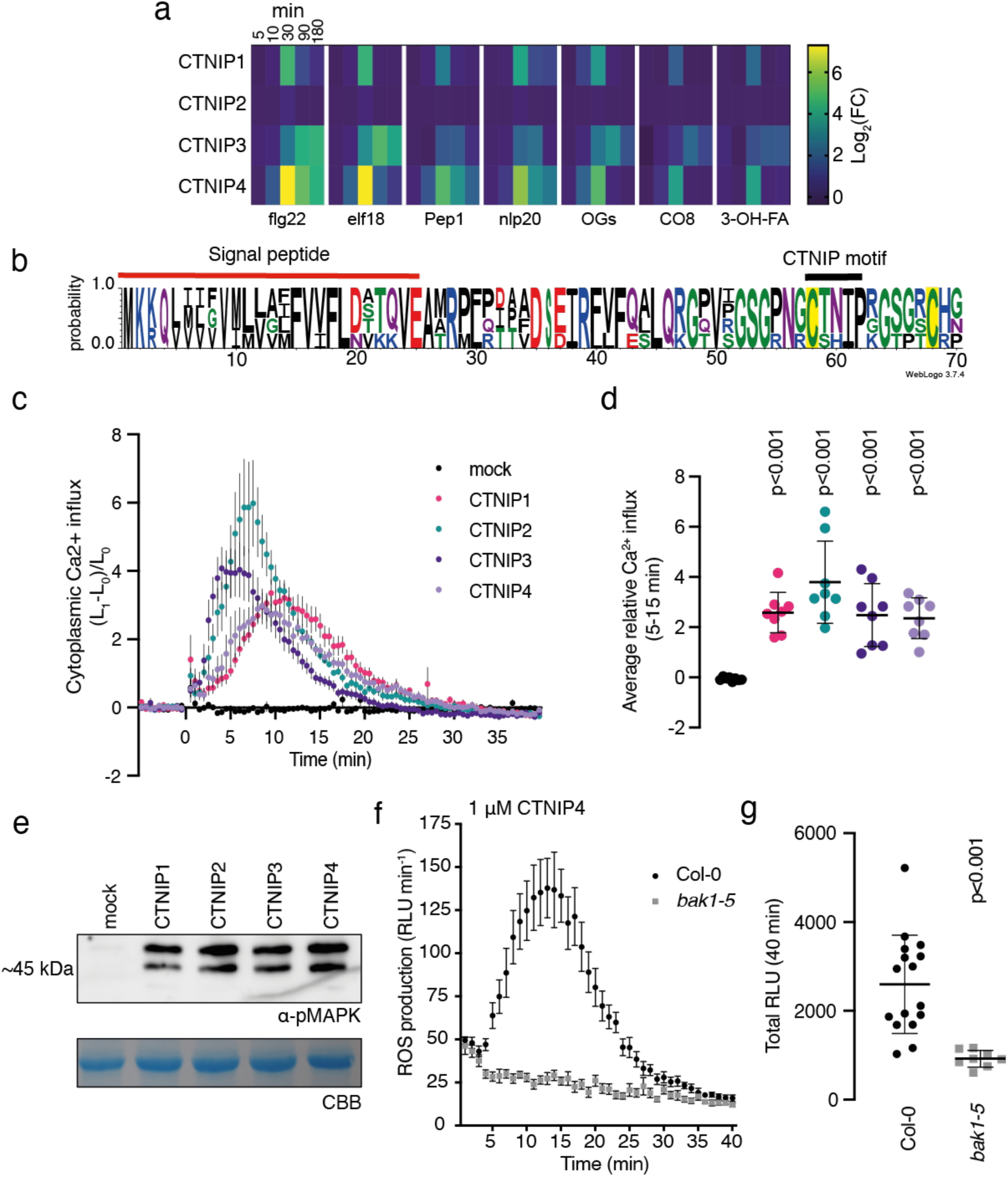
CTNIPs are a novel family of plant signalling peptide. (**a**) Heat map showing log_2_(FC) expression levels of CTNIP1-4 in response to a range of elicitors (data obtained from Bjornson *et al*. (2021)). (**b**) Sequence probability logo from *Arabidopsis* CTNIP1-4 generated using WebLogo3. Signal peptide (as predicted by SignalP5.0) and CTNIP motif are indicated, and conserved cysteine residues are highlighted in yellow. Amino acids are coloured based on their biochemical properties: red = acidic; blue= basic; black = hydrophobic and green = polar. (**c-d**) cytoplasmic calcium influx measured in *p35S::AEQUORIN* seedlings after treatment with 1 μM CTNIP relative to pre-treated levels (*n* = 8 seedlings). (c) Points represent mean; error bars represent S.E.M. (d) represents mean relative Ca^2+^ influx between 5 and 15 min. A line represents mean; error bars represent S.D. *P*-values indicate significance relative to the WT control in a Dunnett’s multiple comparison test following one-way ANOVA. (**e**) Western blot using α-p42/p44-ERK recognizing phosphorylated MAP kinases in seedlings treated with 100 nM CTNIPs or mock for 15 min. The membrane was stained with CBB, as a loading control. (**f-g**) ROS production in leaf disks collected from 4-week-old Arabidopsis plants induced by 1 μM CTNIP4 application (*n* ≥ 8). (f) Points represent mean; error bars represent S.E.M. (g) Integrated ROS production over 40 min. Line represents mean; error bars represent S.D. *P*-values indicate significance relative to the WT control in a two-tailed T-test. All experiments were repeated and analysed three times with similar results. ROS, reactive oxygen species; CBB, Coomassie brilliant blue.

To determine whether CTNIPs function as signalling peptides, we synthetized peptides corresponding to the whole CTNIP proteins excluding the predicted signal peptide. CTNIP1-4 peptides were able to induce cytoplasmic Ca^2+^ influx and mitogen-activated protein kinase (MAPK) phosphorylation – hallmarks of peptide signalling (Figure 1c-e). However, a synthetic peptide derived from the divergent CTNIP5 peptide was inactive (Figure1-figure supplement 1a and 2c). Notably, the C-terminal 23 amino acids of CTNIP4, CTNIP4^48-70^, were sufficient to induce responses (Figure1-figure supplement 2a-b), suggesting that the minimal bioactive peptide is contained within this region. Notably, this region contains two highly conserved cysteine residues (Fig 1b). Mutation of these cysteine residues revealed they are required for CTNIP4 activity (Figure1-figure supplement 2c). Going forward, we focused on CTNIP4 as a representative member of this peptide family, as its transcript was the most up-regulated upon elicitor treatment (Figure 1a).

We hypothesised that CTNIPs may be perceived by a cell-surface LRR-receptor. Typically LRR-receptors are dependent upon the SOMATIC EMBRYOGENESIS RECEPTOR KINASE (SERK) family of co-receptors (Hohmann et al., 2017). We therefore tested whether CTNIP-induced responses were affected in *bak1-5*, an allele of *BRASSINOSTEROID INSENSITIVE 1-ASSOCIATED KINASE 1* (*BAK1/SERK3*) that has a dominant-negative impact on SERK signalling (Perraki et al., 2018; Schwessinger et al., 2011). Concordant with perception by an LRR-receptor, we observed significantly impaired CTNIP4-induced reactive oxygen species production in *bak1-5* (Figure 1f-g).

Ligand-binding induces receptor-SERK heterodimerisation to activate signalling (Hohmann et al., 2017). To identify the CTNIP receptor, we therefore employed Arabidopsis lines expressing BAK1 tagged with green fluorescent protein (GFP) as a molecular bait to identify the CTNIP receptor. Using affinity-purification followed by mass spectrometry we looked for proteins specifically enriched into the BAK1 complex upon CTNIP4 treatment (Figure 2a) (Saur et al., 2016). In four independent biological replicates, the protein most enriched in the BAK1 complex upon CTNIP4 treatment was the LRR-RK HAESA-LIKE 3 (HSL3) (Figure 2b; Figure2-figure supplement 1; Supplementary file 1), making this a promising candidate for being the CTNIP receptor. We could independently confirm CTNIP-induced HSL3-BAK1 complex formation by co-immunoprecipitation (Figure 2c).

**Figure 2.**
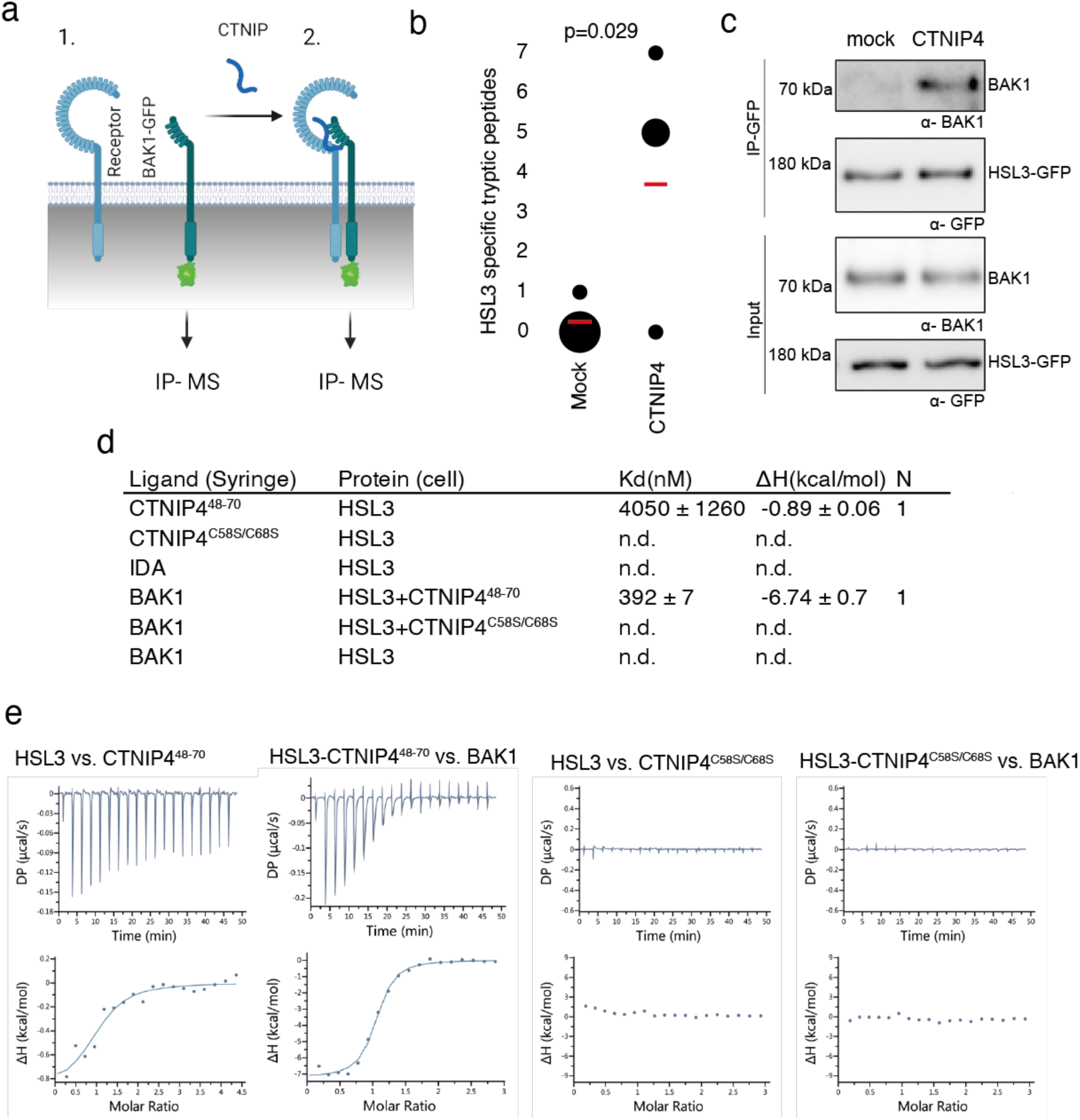
HLS3 forms a CTNIP-induced receptor complex with BAK1. (**a**) Schematic representation of BAK1-GFP immunoprecipitation in the (1) absence or (2) presence of CTNIP4 treatment to identify protein associations induced by CTNIP. Figure generated using Biorender. (**b**) HSL3-specific spectral counts identified in four independent biological replicates where BAK1-GFP was pulled-down in the presence or absence of 1 μM CTNIP4 treatment. Circle diameter is proportional to the number of replicates. Red lines indicate the mean spectral counts for each treatment. *P*-values indicate significance relative to the untreated control in a two-tailed T-test. (**c**) Co-immunoprecipitation of BAK1 with HSL3–GFP from HSL3-GFP seedlings treated with 1 μM CTNIP4^48-70^ or water for 10 min. Western blots were probed with antibodies α-GFP and α-BAK1. This experiment was repeated 3 three times with similar results. (**d**) ITC summary table of HSL3 vs CTNP4^48-70^, CTNP4^C58S/C68S^ and IDA peptides, and contribution of the BAK1 co-receptor to the ternary complex formation. *K_d_*, (dissociation constant) indicates the binding affinity between the two molecules considered (nM). The N indicates the reaction stoichiometry (N=1 for a 1:1 interaction). The values indicated in the table are the mean ± S.E.M. of two independent experiments. (**e**) Isothermal titration calorimetry (ITC) experiments of HSL3 vs CTNIP4 and CTNIP4^C58S/C68S^, in the absence and presence of the co-receptor BAK1.

Consistent with a receptor function, the HSL3 ectodomain (HSL3^ECD^, residues 22-627) heterologously-expressed in insect cells could directly bind CTNIP4 with a dissociation constant of ∼4 µM in *in vitro* binding assays using isothermal titration calorimetry (Figure 2d-e; Figure2-figure supplement 2a-b). In the presence of CTNIP4, BAK1 strongly bound HSL3 with a dissociation constant in the mid-nanomolar range (∼392 nM) (Figure 2d-f; Figure2-figure supplement 2b), consistent with its role as co-receptor. Furthermore, the two conserved cysteine residues are required for receptor binding and co-receptor recruitment explaining their loss of signalling activity (Figure 2d, g-h; Figure1-figure supplement 2c; Figure2-figure supplement 2b).

Notably, we were unable to detect binding of INFLORESCENCE DEFICIENT IN ABSCISSION (IDA), the ligand for the related receptors HAESA and HAESA-LIKE 2 (HSL2) (Meng et al., 2016; Santiago et al., 2016), to HSL3^ECD^ (Figure 2d; Figure2-figure supplement 2b), demonstrating distinct ligand specificity. Accordingly, structural analysis of a HSL3^ECD^ homology model reveals that the HSL3 receptor lacks key conserved motifs required to recognise IDA peptides (Figure2-figure supplement 3) (Santiago et al., 2016). Together, our data show that, while HSL3 is phylogenetically related to HAE, HSL1 and HSL2, it perceives distinct peptides (*i.e.* CTNIPs) most likely via different binding interfaces, which remain to be investigated in future structural studies.

Having established biochemically that HSL3 is the CTNIP receptor, we tested its genetic requirement for CNTIP-induced responses. As expected, we found that HSL3 is strictly required for CTNIP-induced MAPK phosphorylation and whole genome transcriptional reprogramming (Figure 3a-b; Figure3-figure supplement 1). Notably, whilst 30 min treatment with 100 nM CTNIP4 led to differential expression of 1074 genes in wild-type Col-0, none were differentially expressed in *hsl3-1* (p<0.05, |Log2(FC)|>1) (Fig 3b; Supplementary file 2).

**Figure 3.**
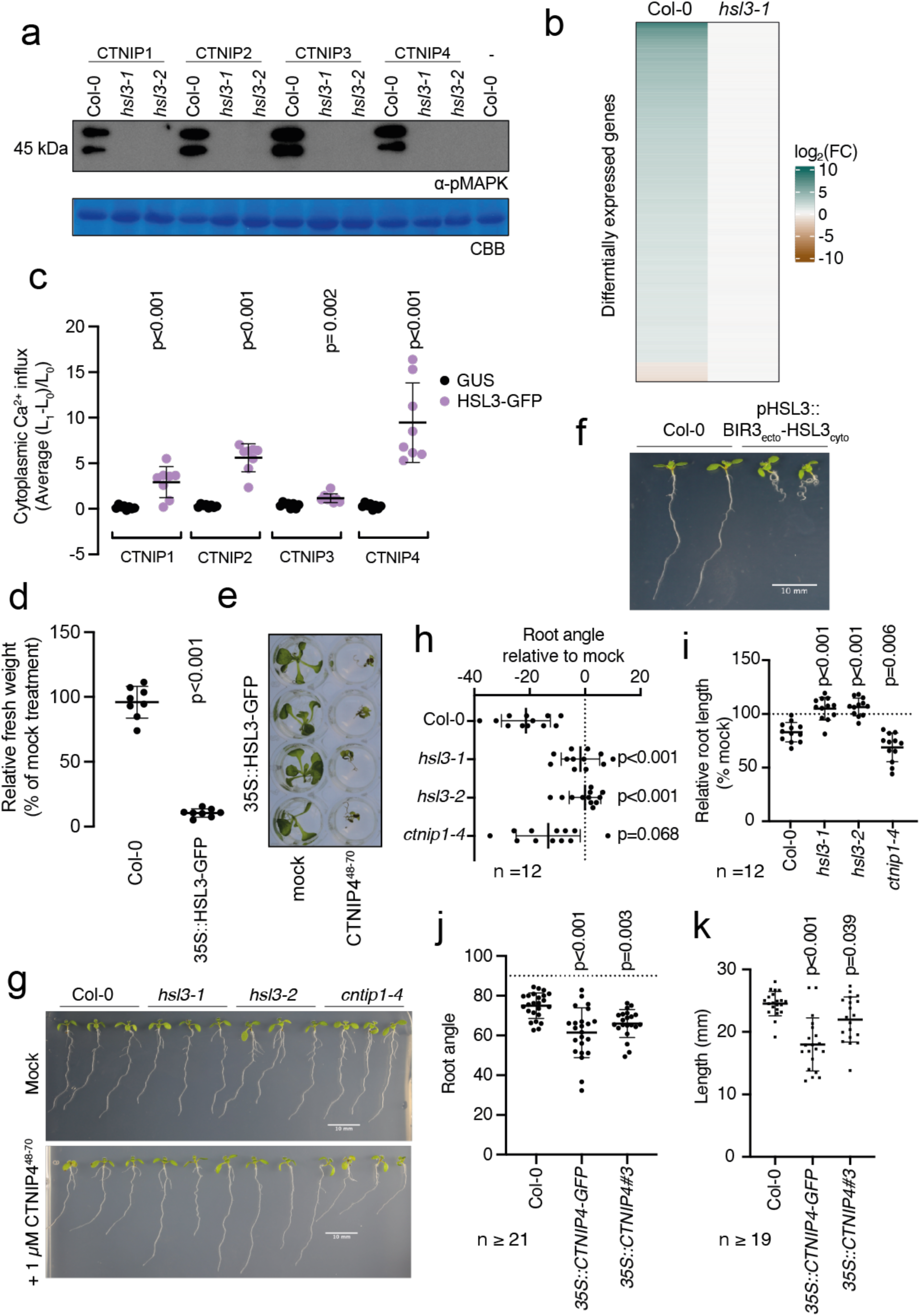
HSL3 is strictly required for CTNIP perception and growth regulation. (**a**) Western blot using α-p42/p44-ERK recognizing phosphorylated MAP kinases in seedlings treated with 100 nM CTNIPs or mock for 15 min. The membrane was stained with CBB, as a loading control. (**b**) Heat map showing all significantly differentially expressed genes (p<0.05, |Log_2_(FC)|>1) in *Arabidopsis* WT or *hsl3-1* seedlings treated with or without 100 nM CTNIP4^48-70^ for 30 min relative to a mock control. (**c**) Mean relative cytoplasmic Ca^2+^ influx in leaf disks of *N. benthamiana* transiently expressing the defined constructs induced by 1 μM CTNIP or mock application (*n* = 8 leaf disks). A line represents mean; error bars represent S.D. *P*-values indicate significance relative to the GUS-transformed control in a Dunnett’s multiple comparison test following one-way ANOVA. (**d**) Fresh weight of 14-day-old seedlings grown in the presence of 500 nM CTNIP4 for 10 days relative to mock (*n* = 8 seedlings). A line represents mean; error bars represent S.D. *P*-values indicate significance relative to the WT control in a two-tailed T-test. (**e**) Representative images of (d). (**f-g**) Nine-day-old vertically grown *Arabidopsis* seedlings on 1/2 MS agar medium with 1 % sucrose. Pictures were taken from the front of the plate. (**h-k**) Root parameters were quantified from the base of the hypocotyl to the root tip using ImageJ (h) Root angle is shown relative to mock. Negative values indicate leftward root skewing. (j) Absolute root angle with 90° representing the gravity vector. Angles <90° represent skewing to the left. A line represents mean; error bars represent S.D. *P*-values indicate significance relative to the WT control in a Dunnett’s multiple comparison test following one-way ANOVA. All experiments were repeated and analysed three times with similar results.

We could additionally show that transient expression of HSL3 in *Nicotiana benthamiana* is sufficient to confer responsiveness to CTNIPs (Figure 3c). Furthermore, whilst 500 nM CTNIP4 was unable to significantly inhibit growth in Col-0 seedlings, plants that over-express *HSL3* became hypersensitive to active CTNIP4 (Figure 3 d-e; Figure3-figure supplement 2). Taken together, our biochemical and genetic results demonstrate that HSL3 is the CTNIP receptor.

CTNIPs induce general early signalling outputs indicative of RK signalling, including cytoplasmic Ca^2+^ influx, MAPK phosphorylation and ROS production (Figure 1) (Olsson et al., 2019). In addition, CTNIP4 treatment induces significant HSL3-dependent transcriptional reprogramming (Figure 3b). Consistent with the up-regulation of *CTNIP* and *HSL3* expression by biotic elicitors (Figure 1a; Figure2-figure supplement 1), gene ontology analysis highlighted the enrichment of many defence- and stress-responsive pathways upon CTNIP4 treatment (Supplementary file 3). This is a pattern shared with other biotic elicitors (Figure3-figure supplement 3) indicative of a general stress response (Bjornson et al., 2021).

To investigate the biological consequence of HSL3 signalling, we fused the extracellular and transmembrane domains of BAK1-INTERACTING RECEPTOR-LIKE KINASE 3 (BIR3) to the cytoplasmic domain of HSL3 under the control of the *HSL3* promoter (Figure3-figure supplement 4a). This chimeric approach allows constitutive complex formation with SERKs, thus mimicking constitutive activation of an endogenous receptor kinase (Hohmann et al., 2020). Transgenic lines expressing this chimeric construct exhibited developmental defects, notably enhanced root curling (Figure 3f). Similarly, CTNIP4 treatment inhibited root growth and induced root skewing in a HSL3-dependent manner (Figure 3 g-i; Figure3-figure supplement 4c). In addition, *CTNIP4* overexpression, either with or without a C-terminal tag, was sufficient to induce a similar phenotype (Figure 3j-k; Figure3-figure supplement 4c). These data suggest that the HSL3-CTNIP signalling module modulates root development, similar to other LRR-RK subfamily XI signalling modules (Jeon et al., 2021; Jourquin et al., 2020).

Recent phylogenetic analyses indicate that HSL3 is conserved in angiosperms (Figure 4a; (Furumizu et al., 2021; Man et al., 2020). Having defined HLS3 as the CTNIP receptor, we wondered whether CTNIPs were equally conserved. CTNIPs were identified in *Amborella*, eudicots and early divergent monocots (Figure 4b-d).

**Figure 4.**
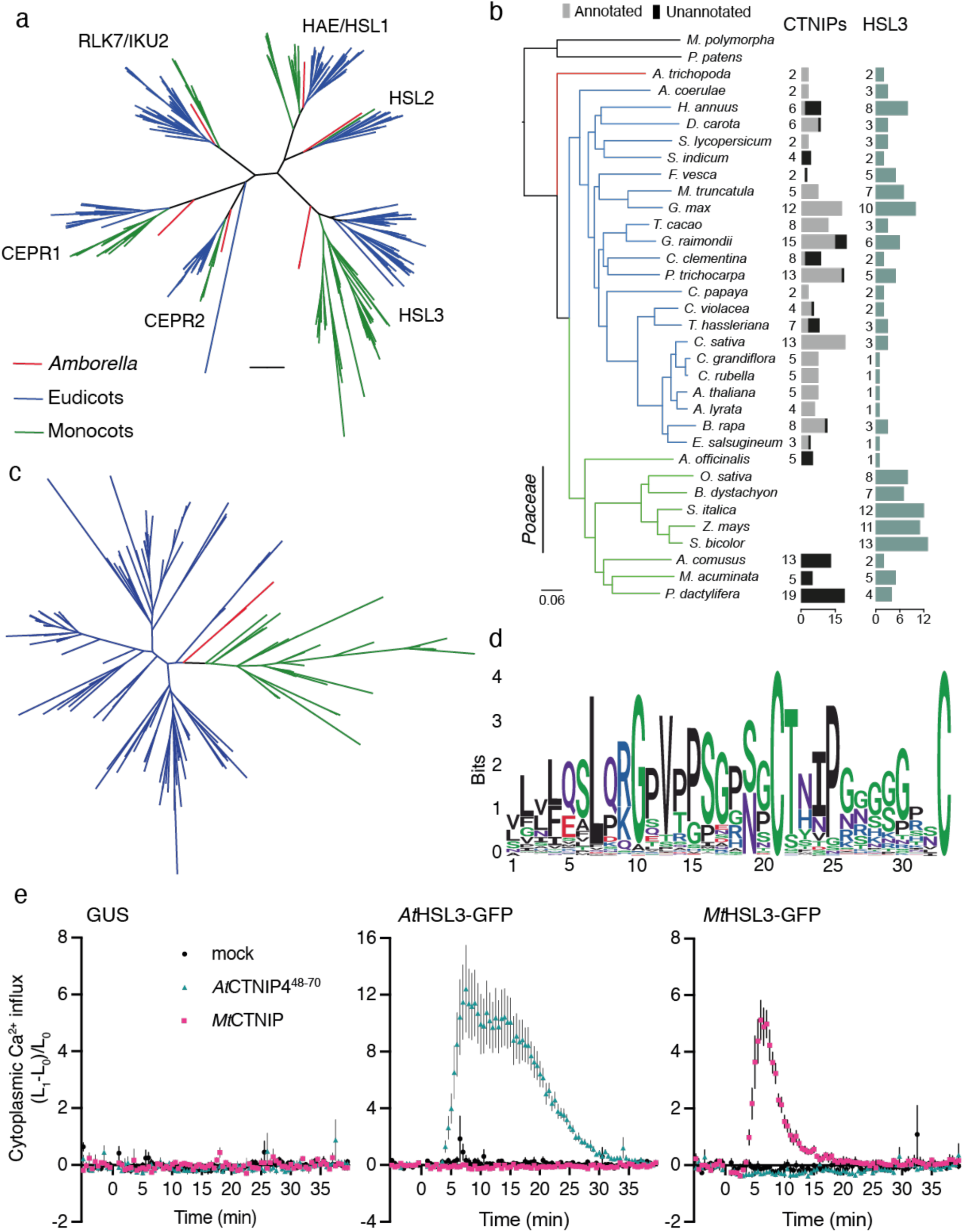
The HSL3-CTNIP signalling module is ancient and conserved. (**a**) Phylogeny of the full-length amino acid sequences of HAE/HSL/CEPR/RLK7/IKU2 clade of receptor kinases. Eudicot sequences are indicated in blue, monocot sequences in green and *Amborella* sequences in red. Clades are named based upon the *Arabidopsis* genes. Alignment shown in Supplementary file 11. Further details of species, sequence identification, alignment and phylogeny generation are described in the material and methods. (**b**) Species tree with number of CTNIP and HSL3 orthologs identified. Annotated CTNIPs are shown in grey whilst unannotated CTNIPs are shown in black. Sequences are shown in Supplementary file 9 and Supplementary file 11. (**c**) Phylogeny of the full-length amino acid sequences of CTNIPs. Eudicot sequences are indicated in blue, monocot sequences in green and *Amborella* sequences in red. Sequences shown in Supplementary file 9. Further details of species, sequence identification, alignment and phylogeny generation are described in the material and methods. (**d**) Sequence Logo generated from CTNIP alignment from (c) using the R-package ggseqlogo. Amino acids are coloured based on their biochemical properties: red = acidic; blue= basic; black = hydrophobic; purple = neutral and green = polar. (**e**) Cytoplasmic calcium influx measured after treatment with 1 μM CTNIP in *p35S::AEQUORIN N. benthamiana* leaf disks transiently expressing the defined construct, relative to pre-treatment (*n* = 8 leaf disks). Points represent mean; error bars represent S.E.M. Experiments were repeated and analysed three times with similar results.

Given the conservation of the HSL3-CTNIP signalling module, the lack of *At*CTNIP4 responses in *N. benthamiana* suggests a co-evolution of ligand-receptor specificity, as for example previously proposed for PLANT ELICITOR PEPTIDE (PEP)-PEP RECEPTOR (PEPR) pairs (Huffaker, 2015; Lori et al., 2015). Accordingly, *Medicago truncatula* HSL3 (*MtHSL3*) only induced a cytoplasmic calcium influx upon treatment with a conspecific CTNIP (*Mt*CTNIP, Medtr1g044470) (Figure 4e).

Our phylogenetic analysis however surprisingly revealed that no clear CTNIP could be found in *Poaceae* genomes (Figure 4b-d). Interestingly, this absence is correlated with an expansion of HSL3 paralogs within these genomes (Figure 4b). We can speculate that the HSL3-CTNIP signalling module may have diverged considerably in this lineage. This is supported by the divergence between eudicot and monocot CTNIPs (Figure 4c). Interestingly, over 40 % of the CTNIPs identified were unannotated, including all monocot CTNIPs (Figure 4b), highlighting how genome annotation still represents a significant challenge in the characterisation of signalling peptides.

## Conclusion

Here, we identified CTNIPs as a novel family of stress-induced signalling peptide. Using affinity-purification and mass spectrometry based on ligand-induced association with the BAK1 co-receptor, we identified the LRR-RK HSL3 as the CTNIP receptor. CTNIPs directly bind the HSL3 ectodomain to promote BAK1 recruitment, and HSL3 is necessary and sufficient to confer CTNIP perception. This ancient signalling module has been conserved for more than 180 million years (Furumizu et al., 2021; Kumar et al., 2017); however, its physiological role remains elusive. HSL3 has recently been shown to play a role in regulating drought and disease resistance implicating HSL3 in multiple stress responses (Lee et al., 2020). Deorphanising HSL3 makes LRR-RK subfamily XI an exciting tool to understand receptor-ligand co-evolution and recognition specificity.

## Acknowledgements

We thank the John Innes Centre Horticultural Services for plant care, especially T. Wells; M. Smoker, J. Taylor and A. Wawryk from the TSL Plant Transformation support group for plant transformation and all past and current members of the Zipfel and Santiago groups for technical help and fruitful discussions. N. Talbot is acknowledged for hosting J.R. for part of this study. This work was supported by the European Research Council under the Grant Agreements no. 773153 and no. 716358 (grant “IMMUNO-PEPTALK” to C.Z. and grant “WallWatchers” to J.S., respectively), The Gatsby Charitable Foundation (to C.Z.), the University of Zürich (to C.Z.), the Swiss National Science Foundation grants no. 31003A_182625 (to C.Z.) and no. 31003A_173101 (to J.S.), and the Fondation Philanthropique Famille Sandoz (to J.S.). M.B. was partially supported by the European Union’s Horizon 2020 Research and Innovation Program under Marie Skłodowska-Curie Actions (grant agreement no. 703954).

---

**Figure 1 – figure supplement 1.**
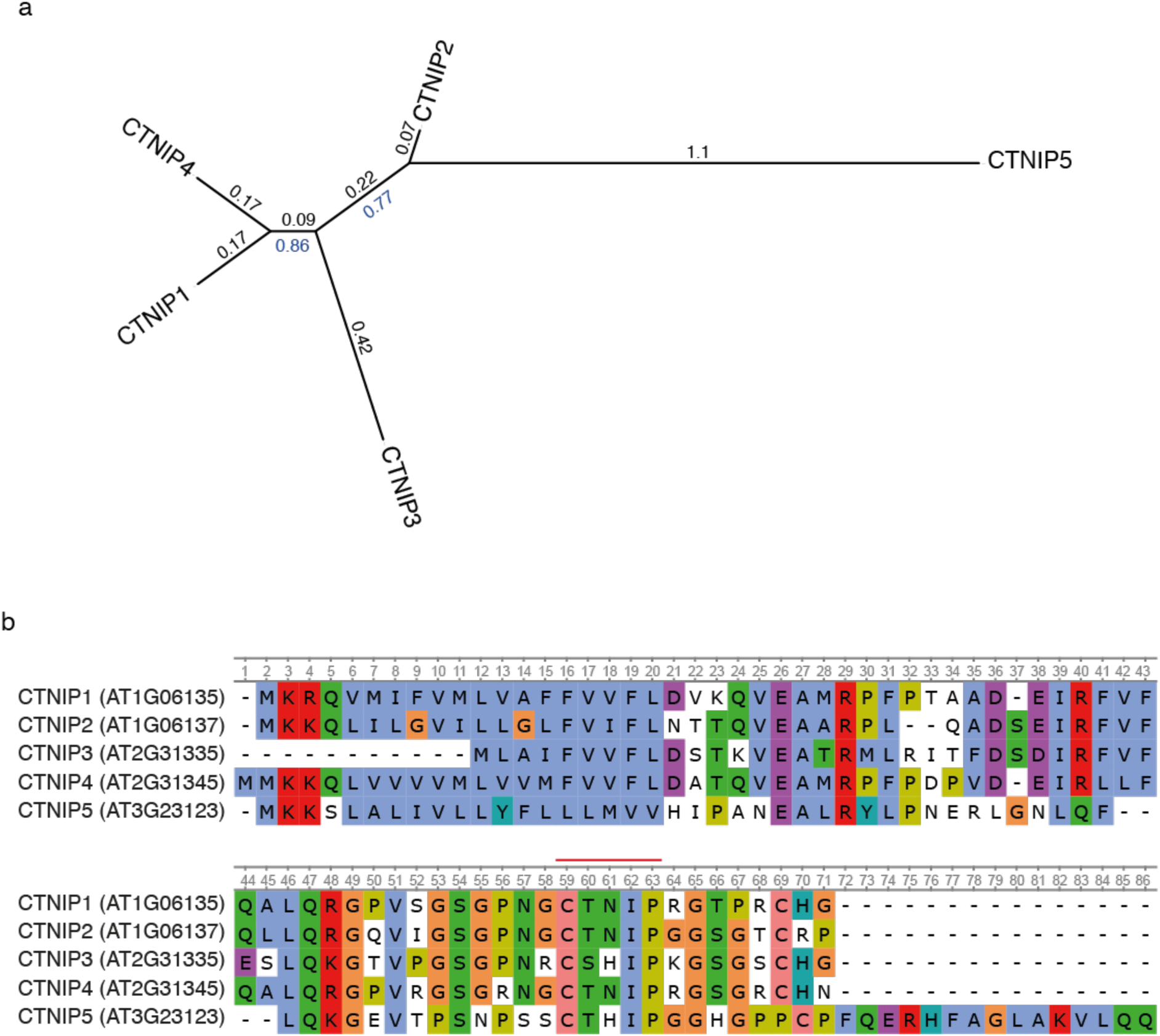
Alignment and phylogeny of *Arabidopsis* CTNIPs. (**a**) Phylogeny of *Arabidopsis* CTNIPs. Full length protein sequences were aligned using MUSCLE and a phylogeny was inferred using the Maximum-likelihood method and JTT matrix-based model conducted in MEGAX. 1000 bootstraps were performed and values shown in blue. Branch lengths are shown in black. (**b**) Alignment used to generate (a). CTNIP motif is highlighted in red.

**Figure 1 -figure supplement 2.**
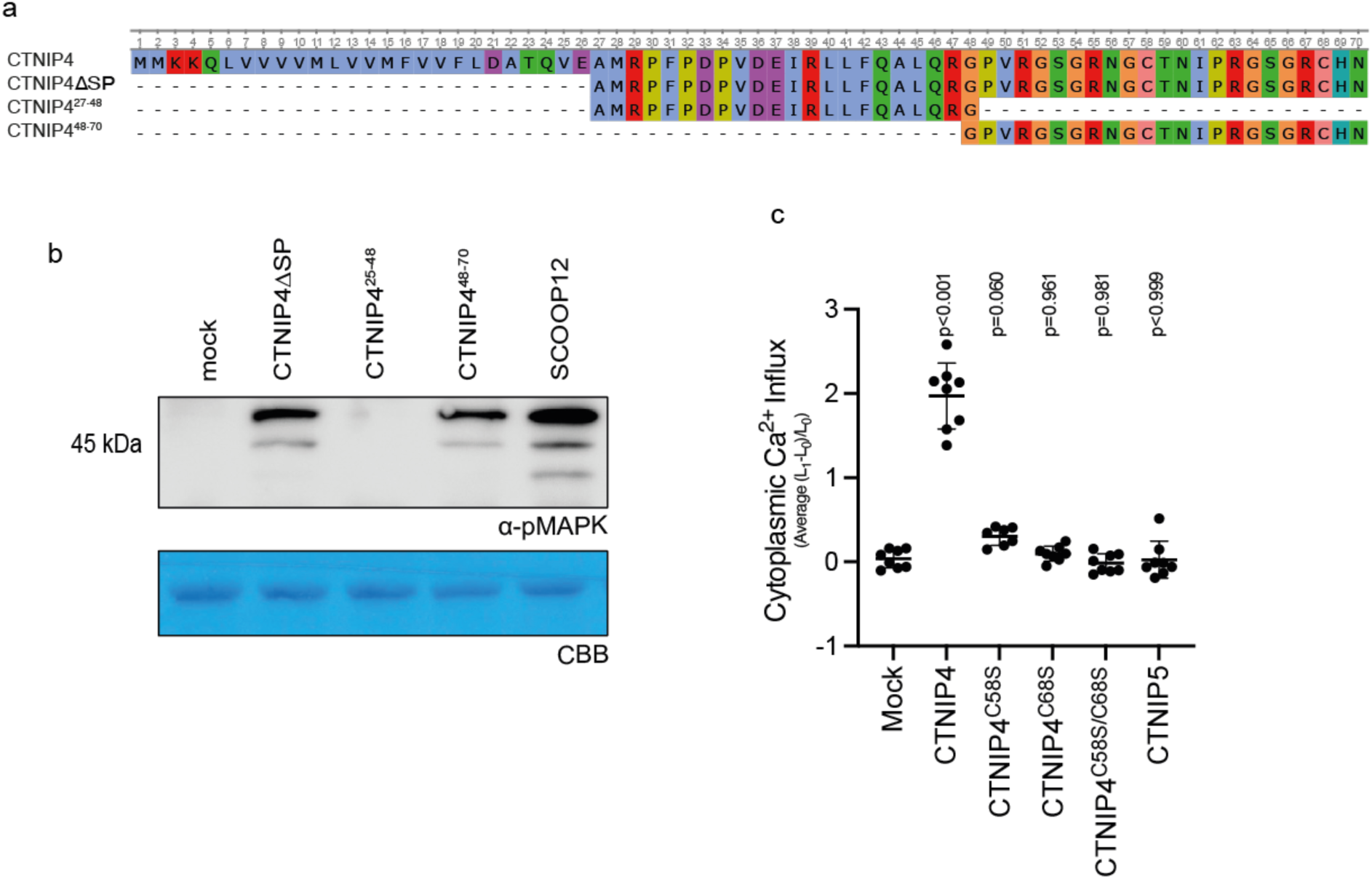
Characterisation of CTNIP fragments. (**a**) Alignment of CTNIP4 fragments used in this manuscript. (**b**) Western blot using α-p42/p44-ERK recognizing phosphorylated MAP kinases in seedlings treated with 100 nM CTNIP4 fragments or mock for 15 min. The membrane was stained with CBB, as a loading control. (**c**) Mean relative Ca^2+^ influx induced by 1 μM CTNIP in *p35S::AEQUORIN* seedlings between 5 and 15 min, relative to pre-treatment (*n* = 8 seedlings). A line represents mean; error bars represent S.D. *P*-values indicate significance relative to the WT control in a Dunnett’s multiple comparison test following one-way ANOVA.

**Figure 2 – figure supplement 1.**
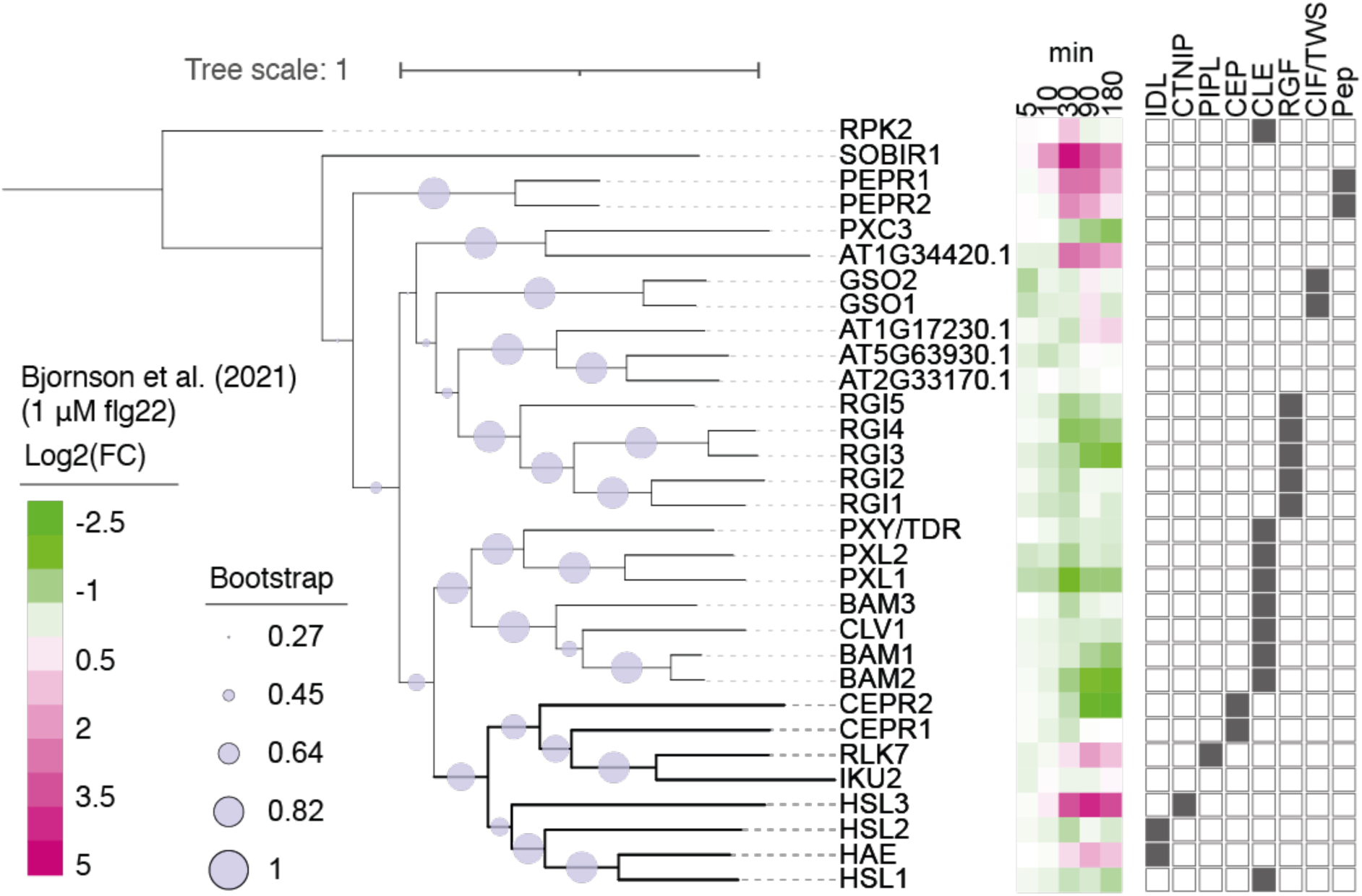
*Arabidopsis* LRR-RK subfamily XI. Phylogeny of full-length protein sequences of the *Arabidopsis* LRR-RK subfamily XI. Sequences were aligned using MUSCLE and a phylogeny was inferred using the Maximum-likelihood method and JTT matrix-based model conducted in MEGAX. 1000 bootstraps were performed and are indicated based on the size of the blue circles. Expression of these receptors in response to 1 μM flg22 treatment was extracted from Bjornson *et al*. (2021) and is represented in a heat map. Known ligands for LRR-RK subfamily XI are highlighted to the right (Butenko et al., 2003; Cho et al., 2008; Crook et al., 2020; Doblas et al., 2017; Doll et al., 2020; Hou et al., 2014; Krol et al., 2010; Morita et al., 2016; Mou et al., 2017; Nakayama et al., 2017; Ogawa et al., 2008; Okuda et al., 2020; Ou et al., 2016; Qian et al., 2018; Rojo et al., 2002; Santiago et al., 2016; Shinohara et al., 2016; Song et al., 2016; Tabata et al., 2014; Yamaguchi et al., 2010, 2006; Zhang et al., 2016).

**Figure 2 – figure supplement 2.**
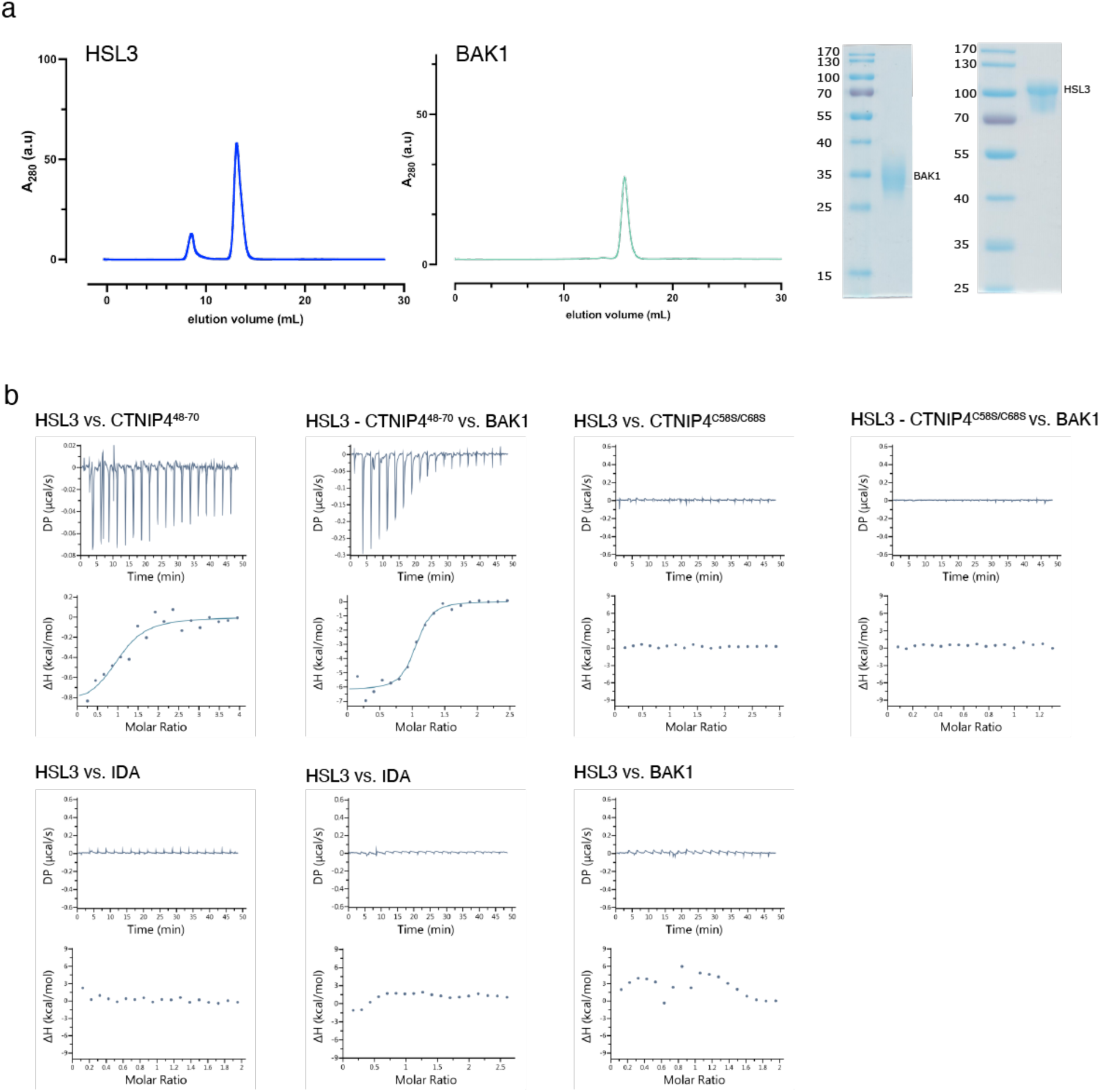
ITC independent experiments and purification of HSL3 and BAK1 used in the binding experiments. (**a**) Analytical size-exclusion chromatography (SEC) of the ectodomains of HSL3 and BAK1. An SDS PAGE of the two proteins in shown alongside. (**b**) ITC raw thermograms of experiments shown in the ITC table summary in Figure 2d.

**Figure 2 – figure supplement 3.**
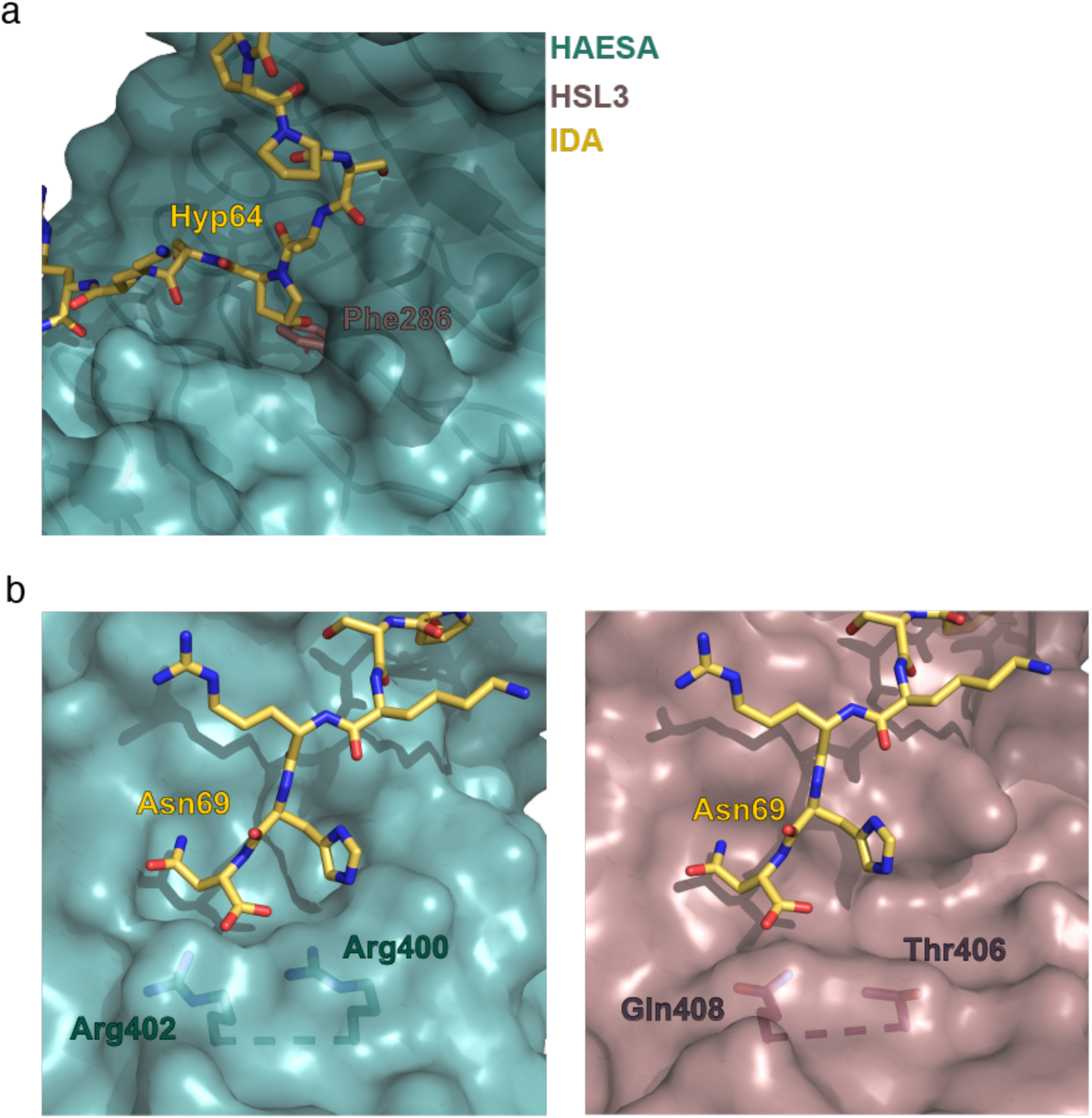
Structural comparison of the binding pockets between the receptors HAESA and HSL3. (**a**) The hydroxyproline pocket required for anchoring the IDA peptide to the HAESA receptor is missing in HSL3. Close view of the binding pocket of the structural superimposition of the HAESA-IDA complex (PDB:5IXQ) and a homology model of HSL3 (AlphaFold: https://alphafold.ebi.ac.uk/). The HAESA receptor is depicted in surface representation in teal blue, IDA in yellow sticks and HSL3 in magenta cartoon. In HSL3, the hydroxyproline pocket is replaced by the bulky residue Phe286, colliding with the potential anchoring of the IDA peptide to the receptor. (**b**) The conserved RxR motif necessary for the coordination of the COO-group the last Asn in IDA is not present in the HSL3 receptor. Zoom in of the C-terminal region of the peptide binding surface of HAESA (teal blue) (left panel) and HSL3 (magenta) (right panel). In HAESA the motif RxR closes the binding pocket allowing for the coordination of the C-terminal of IDA. In HSL3 this structural motif is substituted by the residues Thr406 and Gln408, leaving the binding surface open to potentially accommodate a longer peptide ligand. Figures were done using the PyMOL Molecular Graphics System, Version 2.0 Schrödinger, LLC.

**Figure 3 – figure supplement 1.**
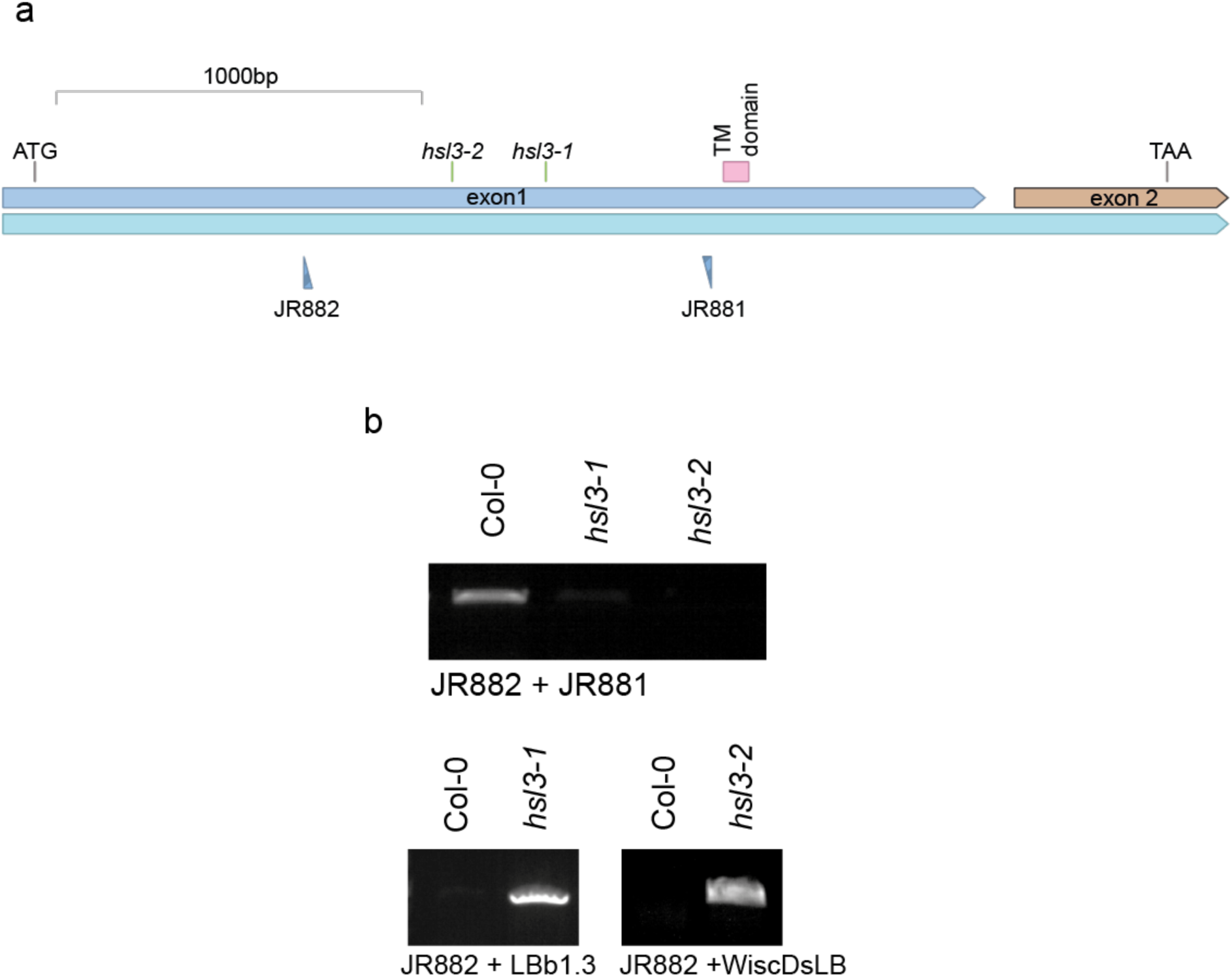
Genetic characterization of *hsl3* mutants. (**a**) Gene model showing the location of T-DNA inserts. (**b**) PCR confirming T-DNA insertion and mutant homozygosity.

**Figure 3 – figure supplement 2.**
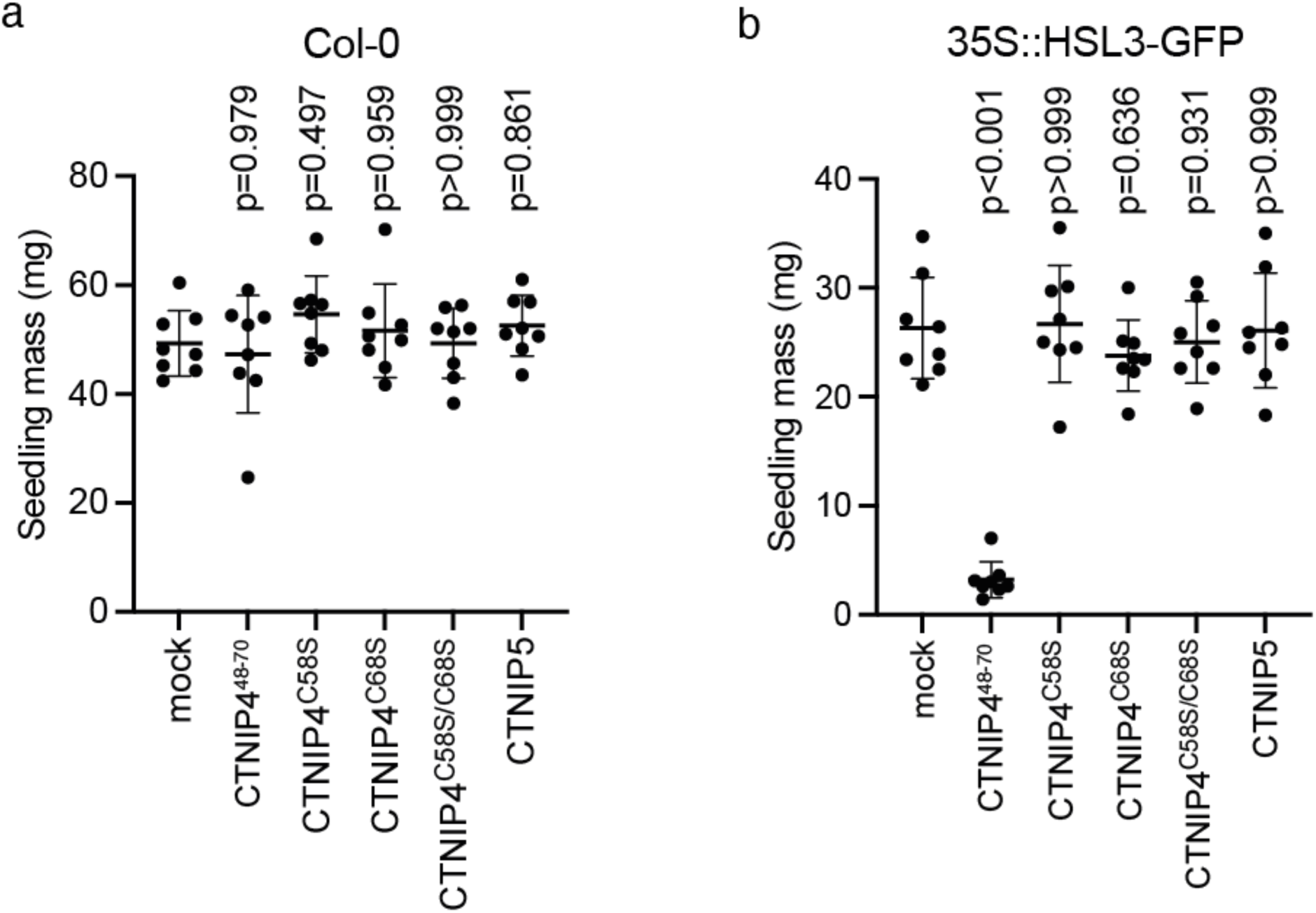
CTNIP-induced seedling growth inhibition. (**a-b**) Fresh weight of 14-day-old seedlings grown in the presence of 500 nM CTNIPs for 10 days relative to mock (*n* = 8 seedlings). A line represents mean; error bars represent S.D.; *P*-values indicate significance relative to the WT control in a Dunnett’s multiple comparison test following one-way ANOVA.

**Figure 3 – figure supplement 3.**
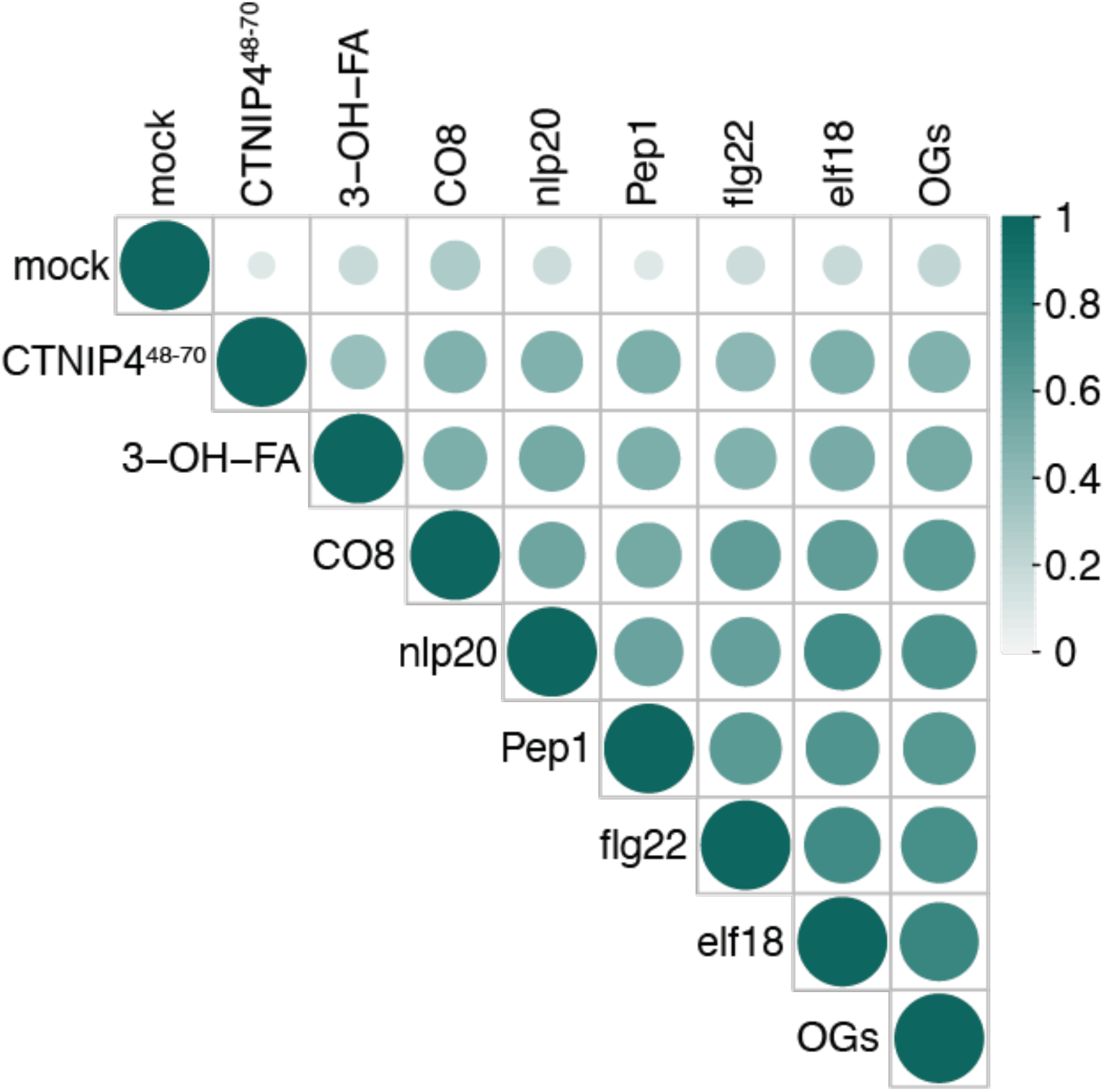
Correlation of CTNIP4-induced transcriptomic response with that of elicitors at 30 min. CTINP4-induced gene expression is well correlated with elicitor-induced gene expression from Bjornson *et al*. (2021). Circle colour and size are proportional to the Spearman correlation coefficient (R-squared value) of each pairwise comparison of log_2_(fold changes).

**Figure 3 – figure 4.**
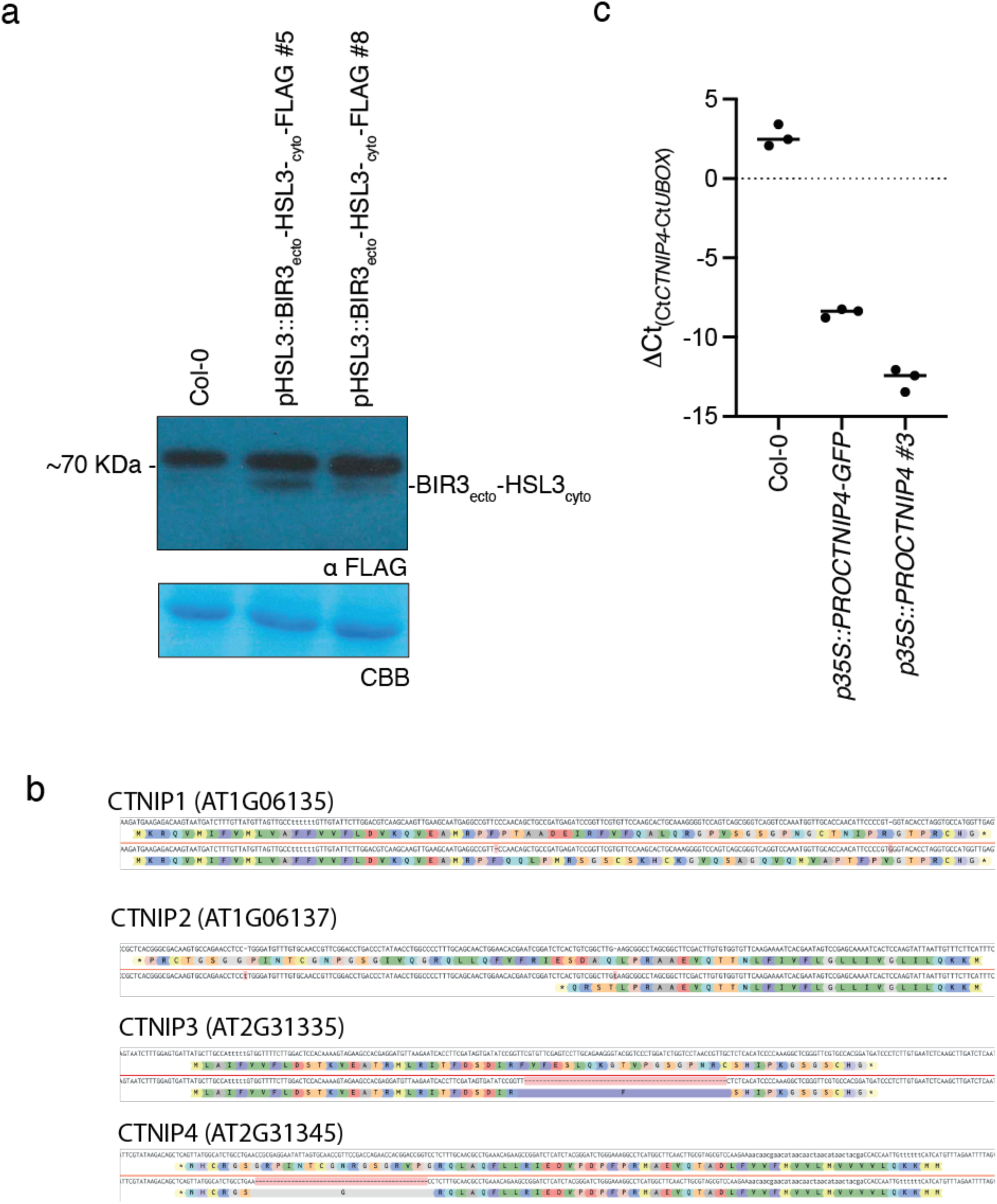
Characterisation of CTNIP and chimeric receptor lines. (**a**) Western blot using α-FLAG recognizing BIR3_ecto_-HSL3_cyto_-FLAG in seedlings to confirm expression. The membrane was stained with CBB, as a loading control. (**b**) Documentation of the Cas9-induced mutations observed within the *ctnip1-4* polymutant and their predicted effects on protein products. (**c**) qRT-PCR documenting the overexpression of CTNIP lines. Expression of *CTNIP4* is shown relative to *U-Box* (*At5g15400*). Points represent independent biological replicates each comprising two technical replicates. Lines represent the mean of biological replicates.

**Supplementary file 1. Spectral counts of peptides identified through affinity-purification of the BAK1 complex**

**Supplementary file 2. Differential gene expression induced by 30 min CTNIP4^48-70^ treatment**

**Supplementary file 3. Gene ontology enrichment following 30 min CTNIP4^48-70^ treatment**

**Supplementary file 4. Primers**

**Supplementary file 5. Synthetic peptides**

**Supplementary file 6. Species induced in CTNIP and RK search**

**Supplementary file 7. Initial CTNIP candidates used to search**

**Supplementary file 8. Identified CTNIPs relaxed**

**Supplementary file 9. Identified CTNIPs confident**

**Supplementary file 10. Initial RK candidates used to search**

**Supplementary file 11. Alignment of RKs identified**

## Material and Methods

### Plant material and growth conditions

Arabidopsis plants for ROS burst assays were grown in individual pots at 21 °C with a 10-h photoperiod. Seeds grown on plates were surface sterilized using chlorine gas for 5–6 h and sown on 1/2 Murashige and Skoog (MS) media, 1 % sucrose, and 0.8 % agar and stratified at 4 °C for 2–3 days. Plates were then transferred to 22 °C under a 16-h photoperiod. For root growth assays plates were placed in an upright position under a 10° angle relative to the direction of gravity and images were taken 9 days later (Van der Does et al., 2017). *Nicotiana benthamiana* plants were grown on peat-based media at 24 °C, with 16-h photoperiod.

Aequorin lines of *Arabidopsis* and *N. benthamiana* were described previously (Knight et al., 1991; Segonzac et al., 2011). *hsl3* mutants have been previously described and were obtained from the Eurasian Arabidopsis Stock Centre (uNASC) (Hou et al., 2014; Lee et al., 2020). *bak1-4/*pBAK1::BAK1-GFP, *bak1-5* and p35S::GFP-Lti6B lines have also been described previously (Cutler et al., 2000; Ntoukakis et al., 2011; Schwessinger et al., 2011).

### Synthetic peptides

All synthetic peptides were ordered at >80 % purity from either EZbiolabs or Genscript. Sequences of all peptides can be found in Supplementary file 5.

### Alignment and phylogeny of Arabidopsis CTNIPs and LRR-RK subfamily XI

Full length protein sequences were aligned using MUSCLE (Edgar, 2004) and a phylogeny was inferred using the Maximum-likelihood method and JTT matrix-based model conducted in MEGAX (Kumar et al., 2018). 1000 bootstraps were performed. Trees were visualised in iTOL (Letunic and Bork, 2019). The sequence logo was generated using WebLogo3 (Crooks et al., 2004).

### Molecular cloning

#### In-planta expression

For overexpression of *At*HSL3-GFP and *MtHSL3* in *N. benthamiana* and Arabidopsis the genomic DNA sequence was amplified from Arabidopsis ecotype Columbia and *M. truncatula* ecotype A11, domesticated and directly ligated into pICSL86977 downstream of a 35S promoter and with an in-frame C-terminal GFP tag.

Fragments for the pHSL3::BIR3ecto-HSL3cyto-FLAG construct were amplified from genomic DNA using the indicated primers and ligated into pICSL86955 (Supplementary file 4). Fragments were designed according to Hohmann *et al*. (2020). Clones were verified by Sanger sequencing.

### CRISPR-Cas9 mutagenesis

CRISPR-Cas9 induced mutagenesis was performed as described by Castel *et al*. (2019). The *RPS5a* promoter drove Cas9 expression and FASTred selection was used for positive and negative selection. Primers used to generate the vector can be found in Supplementary file 4. Mutants were screened by Sanger sequencing.

### ROS measurements

Leaf disks were harvested from 4-week-old *Arabidopsis* plants into white 96-well-plates (655075, Greiner Bio-One) containing 100 μL water using a 4-mm diameter biopsy punch (Integra™ Miltex™). Leaf disks were rested overnight. Prior to ROS measurement, the water was removed and replaced with ROS assay solution (100 μM Luminol (123072, Merck), 20 μg mL^−1^ horseradish peroxidase (P6782, Merck)) with or without elicitors. Immediately after light emission was measured from the plate using a HIGH-RESOLUTION PHOTON COUNTING SYSTEM (HRPCS218, Photek) equipped with a 20 mm F1.8 EX DG ASPHERICAL RF WIDE LENS (Sigma Corp).

### Cytoplasmic calcium measurements

Seedlings were initially grown on 1/2 MS plates for 3 days before being transferred to 96-well plates (655075, Greiner Bio-One) in 100 μL liquid MS for 5 days. The evening before calcium measurements the liquid MS was replaced with 100 μL 20 μM coelenterazine (EC14031, Carbosynth) and the seedlings incubated in the dark overnight. The following morning the coelenterazine solution was replaced with 100 μL water and rested for a minimum of 30 min in the dark. Readings were taken in a VARIOSKAN^TM^ MUTIPLATE READER (ThermoFisher) before and after adding 50 μL of 3× concentrated elicitor solution or mock. For each well readings were normalised to the average RLU value before elicitor addition (L_0_).

### Seedling growth inhibition

Four-day-old seedlings growing on 1/2 MS plates were transferred into individual wells of a transparent 48-well tissue culture plate (Greiner Bio-One) containing 500 μL of liquid MS media with/without elicitor addition. The plates were returned to the growth conditions for an additional 10 days before seedlings were blot-dried and weighed.

### Protein extraction and western blot

Two-week-old seedlings grown in liquid MS media (MAPK phosphorylation) or leaf disks from 4-week-old plants were flash-frozen in liquid nitrogen. Frozen plant tissue was ground in a Genogrinder® with 2mm glass beads (1500 strokes/min, 1.5 min) prior to boiling in 2× Laemmli sample buffer (4 % SDS, 20 % glycerol, 10 % 2-mercaptoethanol, 0.004 % bromophenol blue, and 0.125 M Tris-HCl; (10 μL.mg^−1^ tissue)) for 10 min at 95 °C. The samples were then spun at 13,000 rcf for 5 min prior to loading and running on SDS-PAGE gels. Proteins were transferred using semi-dry transfer onto PVDF membrane (ThermoFisher), blocked in 5 % (w/v) Bovine serum albumin prior to incubation with appropriate antibodies (α-pMAPK ((p44/42 MAPK (Erk1/2) antibody #9102; 1:4000); α-FLAG-HRP (A8592, Merck; 1:5,000) and α-rabbit-HRP (A-0545, Merck; 1:10000). Western blots were imaged with a LAS 4000 IMAGEQUANT SYSTEM (GE Healthcare) or on X-ray film before being developed. Staining of the blotted membrane with Coomassie brilliant blue was used to confirm loading.

### Co-immunoprecipitation

All steps involving the protein extract and subsequent protein isolation were carried out on ice or at 4 °C and all buffers and tubes were pre-cooled.

Seeds were sown on 1/2 MS agar and stratified for 3 days as described above. When seedlings had germinated, they were transferred 6 seedlings per well into 6 well plates containing 5 mL of liquid MS media and grown for a further 12 days. Seedlings were then transferred into MS media either with or without CTNIP4 addition and vaccum infiltrated for 2 min and left in the solution for a further 10 min. Seedlings were rapidly dried and flash frozen in liquid nitrogen and ground. Proteins were extracted using by addition of ∼2:1extraction buffer (50 mM Tris pH 7.5, 150 mM NaCl, 2.5 mM EDTA, 10 % Glycerol, 1 % IGEPAL, 5 mM DTT, 1 % plant protease inhibitor cocktail (P9599, Sigma)):ground tissue (v/v). Proteins were solubilised at 4 °C with gentle agitation for 30 min before filtering through miracloth. The filtrate was centrifuged at 30000 rcf for 30 min at 4 °C. Protein concentrations were normalised using Bradford assay. An input sample was taken. To each 15 ml of protein extract 40 μl of GFP-TRAP AGAROSE BEADS (50 % slurry, ChromoTek) washed in extraction buffer were added and incubated with gentle agitation for 4 h at 4 °C. Bead were harvested by centrifugation at 1500 x g for 2 min and washed 3 times in extraction buffer. Beads were then resuspended in fifty microliters of 1.5 x elution (NuPage) buffer and incubated at 80 °C for 8 min. Samples were subsequently used from MS analysis or Western blotting.

Western blotting was performed as described previously (α-BAK1 (Roux et al., 2011); 1:2000), α-GFP-HRP (sc-9996, Santa Cruz; 1:5000) and α-rabbit-HRP (A-0545, Merck; 1:10000).

### Sample Preparation for Mass Spectrometry

Co-immunoprecipitated protein samples were ran approximately 1 cm into an SDS-PAGE gel. This portion of the gel was then excised, cut into smaller pieces and washed three times with acetonitrile (LC-MS-Grade):ammonium bicarbonate (50 mM), pH 8.0 (1:1, v/v), 30 min each, followed by dehydration in acetonitrile, 10 min. Gel pieces were then reduced with 10 mm DTT for 30 min at 45 °C followed by alkylation with 55 mm iodoacetamide for 20 min at room temperature, and a further three washes with acetonitrile:ammonium, 30 min each. Gel pieces were dehydrated again with acetonitrile before rehydration with 40 µL trypsin (Pierce Trypsin Protease, MS-Grade, catalog no. 90058) working solution (100 ng trypsin in 50 mM ammonium bicarbonate, 5% (v/v) acetonitrile). Where required, gel pieces were covered with 50 mM ammonium bicarbonate to a final volume before incubation at 37 °C overnight. Tryptic peptides were extracted from the gel pieces three times in an equal volume of 50% acetonitrile, 5% formic acid (Pierce LC-MS-Grade, catalog no. 85178), 30 min each. Extracted peptides were dried in a speed-vac and resuspended in (v/v) 2% acetonitrile/0.2% trifluoroacetic acid (Merck, catalog no. 302031). A total of four biological replicates for each sample type was submitted.

### LC-MS/MS Analysis

Approximately 35% of each sample was analysed using an Orbitrap Fusion™ Tribrid™ Mass Spectrometer (Thermo Fisher Scientific) coupled to a U3000 nano-UPLC (Thermo Fisher Scientific). The dissolved peptides were injected onto a reverse phase trap column NanoEase m/z Symmetry C18, beads diameter 5 μm, inner diameter 180 μm x 20 mm length (Waters). The column was operated at the flowrate 20μl/min in 2% acetonitrile, 0.05% TFA, after 2.5min the trap column was connected to the analytical column NanoEase m/z HSS C18 T3 Column, beads diameter 1.8 μm, inner diameter 75 μm x 250 mm length (Waters). The column was equilibrated with 3% B (B: 80% acetonitrile in 0.05% formic acid (FA), A: 0.1% FA) before subsequent elution with the following steps of a linear gradient: 2.5min 3% B, 5min 6.3% B, 13min 12.5% B, 50min 42.5% B, 58min 50% B, 61min 65% B, 63min 99% B, 66min 99% B, 67min 3% B, 90min 3% B. The flow rate was set to 200 nL/min. The mass spectrometer was operated in positive ion mode with nano-electrospray ion source. Molecular ions were generated by applying voltage +2.2kV to a conductive union coupling the column outlet with fused silica PicoTip emitter, ID 10 μm (New Objective, Inc.) and the ion transfer capillary temperature was set to 275°C. The mass spectrometer was operated in data-dependent mode using a full scan, m/z range 300–1,800, nominal resolution of 120,000, target value 1 × 10^6^, followed by MS/MS scans of the 40 most abundant ions. MS/MS spectra were acquired using normalized collision energy of 30%, isolation width of 1.6 m/z, resolution of 120,000, and a target value set to 1 × 10^5^. Precursor ions with charge states 2-7 were selected for fragmentation and put on a dynamic exclusion list for 30 seconds. The minimum automatic gain control target was set to 5 × 10^3^ and intensity threshold was calculated to be 4.8 × 10^4^. The peptide match feature was set to the preferred mode and the feature to exclude isotopes was enabled.

### Data Processing and Peptide Identification

Peak lists in the form of Mascot generic files were prepared from raw data files using MS Convert (Proteowizard) and sent to a peptide search on Mascot server v2.7 using Mascot Daemon (Matrix Science, Ltd.) against an in-house contaminants database and the Araport 11 protein database. Tryptic peptides with up to 1 possible mis-cleavage and charge states +2, +3 were allowed in the search. The following peptide modifications were included in the search: carbamidomethylated Cysteine (fixed) and oxidized Methionine (variable). Data were searched with a monoisotopic precursor and fragment ion mass tolerance 10 ppm and 0.8 Da respectively. Decoy database was used to validate peptide sequence matches.

Mascot results were combined in Scaffold v4.4.0 (Proteome Software Inc.) and peptide and protein identifications accepted if peptide probability and protein threshold was ≥ 80.0% and 99% respectively. Under these conditions the False Discovery Rate was 0.47%. Data was then exported to Excel (Microsoft) for further processing. Proteins were accepted if identified by at least 2 peptides and present in 2 or more biological replicates. Spectral counts from the 4 biological replicates were summed and used to derive a ratio of CTNIP4 treatment:mock treatment. The mass spectrometry proteomics data have been deposited to the ProteomeXchange Consortium via the PRIDE (Perez-Riverol et al., 2016) partner repository with the dataset identifier PXD029264 and 10.6019/PXD029264

### Transient expression in *Nicotiana benthamiana*

*Agrobacterium tumefaciens* strain GV3101 transformed with the appropriate construct were grown overnight in L-media and spun-down. The bacteria were resuspended in 10 mM MgCl_2_ and adjusted to O.D._600_ = 0.2 prior to infiltration into the youngest fully expanded leaves of 3-week-old plants. Leaf disks were collected 24 h later, and calcium assays were performed as described for seedlings with leaf disks being floated overnight in the dark in 20 μM coelenterazine (EC14031, Carbosynth).

### Protein expression and purification

The ectodomains expressed and purified were coded from Arabidopsis genes *HSL3* (22-627, AT5G25930) and *BAK1* (residues 20 – 637, AT4G33430). Codon-optimised synthetic genes were cloned into a modified pFastBac vector (Geneva Biotech) vector, providing a TEV (tobacco etch virus protease) cleavable C-terminal StrepII-9xHis tag. Expression of HSL3 and BAK1 was driven by the signal peptides 30K (Futatsumori-Sugai and Tsumoto, 2010) or *Drosophila BiP* (Smakowska-Luzan et al., 2018), respectively. The baculovirus were generated in DH10 cells and *Spodoptera frugiperda* Sf9 cells were used for viral amplification. For protein expression *Trichoplusia ni* Tnao38 cells (Hashimoto et al., 2012), were infected with HSL3 and BAK1 viruses with a multiplicity of infection (MOI) of 3. The cells were grown 1 day at 28 °C and two days at 22 °C at 110 rpm The secreted proteins were purified separately by sequential Ni^2+^ (HisTrap excel, GE Healthcare, equilibrated in 25 mM KP_i_ pH 7.8 and 500 mM NaCl) and StrepII (Strep-Tactin Superflow high-capacity, (IBA, Germany) equilibrated in 25 mM Tris pH 8.0, 250 mM NaCl, 1 mM EDTA) affinity chromatography. Recombinant Strep-tagged TEV protease was used in 1:50 ratio to remove the affinity tags. The cleaved tag and the protease were separated from the protein ectodomains by Ni^2+^ affinity chromatography. Proteins were further purified by size exclusion chromatography on a Superdex 200 Increase 10/300 GL column (GE Healthcare) equilibrated in 20 mM citric acid pH 5.0, 150 mM NaCl. Peak fractions containing the complex were concentrated using Amicon Ultra concentrators (Millipore, MWCO 10,000 for BAK1 and 30,000 for HSL3). Proteins were analysed for purity and structural integrity by SDS-PAGE.

### Isothermal titration calorimetry (ITC)

A MicroCal PEAQ-ITC (Malvern Instruments) was used to performing the ITC binding assays. Experiments were performed at 25 °C with a 200 µL standard cell and a 40 μL titration syringe. HSL3 and BAK1 proteins were gel-filtrated into pH 5 ITC buffer (20 mM sodium citrate pH 5.0, 150 mM NaCl). Protein concentrations for HSL3 and BAK1 were calculated using their molar extinction coefficient and a calculated molecular weight of ∼75,000 for HSL3 and ∼ 25,000 Da for BAK1. Experiments were performed with 20µM of HSL3 protein in the cell and between 200-450 μM of indicated peptide ligand in the syringe, following an injection pattern of 2 μL at 150 s intervals and 500 r.p.m. stirring speed. The BAK1 vs HSL3-peptide experiments were performed by titrating 100 µM of BAK1 in the cell, using the same injection pattern. ITC data were corrected for the heat of dilution by subtracting the mixing enthalpies for titrant solution injections into protein free ITC buffer. Experiments were done in replicates and data were analyzed using the MicroCal PEAQ-ITC Analysis Software provided by the manufacturer. All ITC runs used for data analysis had an N ranging from 0.7 to 1.3. The N values were fitted to 1 in the analysis.

### RNA sequencing and qRT-PCR

Two 3-day-old seedlings per well were transferred into transparent 24-well plates (Grenier Bio-One) containing 1 mL liquid MS media, sealed with porous tape and grown for a further 9 days. For qRT-PCR seedlings were harvested at this point. For RNA-seq experiments media was then exchanged for 500 µL fresh MS media and left overnight. In the morning a further 480 µL of fresh media was added. 9.5 h later 20 µL treatment/mock was added and seedlings were harvested after 30 min. All seedlings were ground in liquid nitrogen.

Total RNA was extracted using Trizol reagent (Merck) according to the manufacturer’s instructions and DNAase/RNA cleanup treatment was performed using the RNeasy kit (Qiagen). RNA sequencing was performed by Novogene. The RNA-seq datasets generated and analysed in the current study have been deposited in the ArrayExpress database at EMBL-EBI (www.ebi.ac.uk/arrayexpress) under accession number E-MTAB-11093. qRT-PCR was performed on cDNA synthesised using The RevertAid first strand cDNA synthesis kit (Thermofisher) according to the manufacturer’s instructions. cDNA was amplified by quantitative PCR using SYBR Green JumpStart Taq ReadyMix (Roche) and the CFX96 Real-Time PCR Detection System (Bio-Rad Laboratories, Hercules, CA, USA).

The read data were analysed using FastQC, trimmed using trimmomatic (Bolger et al., 2014) and mapped to the *Arabidopsis* TAIR10 genome via TopHat2 (Andrews et al., 2015; Kim et al., 2013). The mapped reads were assigned to genes by featureCounts from package Rsubread in R (Liao et al., 2019), and differential expression analysis was performed using DESeq2 with ashr L2FC shrinkage (Love et al., 2014; Stephens, 2017). Changes in gene expression were visualised using the R package ComplexHeatmap (Gu et al., 2016).

### GO enrichment

GO term enrichment was calculated using the R package topGO (Alexa and Rahnenfuhrer, 2021), with arguments method= weight.01 and statistic=Fisher.

### Correlation of expression

Pairwise comparisons of gene expression differences (log_2_(FC)) was performed in R using the rcorr function from package Hmisc (Harrell Jr, 2021), type=Spearman, and correlations were plotted using corrplot (Wei and Simko, 2021).

### Genome data retrieval

Whole genome sequences and protein sequences were retrieved from Ensembl (release 50), Phytozome (version 13), NCBI, and marchantia.info. Species and individual assembly versions are listed in Supplementary file 6 (SI_table_species_data.csv).

### CTNIP identification

#### Peptide search

Protein sequences from all species were first filtered for a maximum length of 300 amino acids and merged into a single file. The initial set of CTNIP peptide sequences is given in Supplementary file 7 (SI_data_initial_CTNIP_candidates.fasta). Additional candidates were searched with 1) jackhmmer (version 3.1b2, (Eddy, 2011)), 2) diamond (version 0.9.26, options -e 1e-8 -k 100, (Buchfink et al., 2014)), and 3) hmm profile search (3.1b2, (Wheeler and Eddy, 2013)). For the hmm profile search, the initial set of candidates and the candidates from the diamond search were aligned with muscle (v3.8.31, (Edgar, 2004)) to generate an hmm profile (hmmbuild) that was then used to search more candidates (hmmsearch). Candidates from all approaches were merged and grouped with a sequence similarity network. For this, sequences were matched to each other with diamond (options -e 0.01 -k 100). The pairwise percent similarity scores above 20 % were used to construct a network. The community structure of the network was resolved with a modularity optimization algorithm (Blondel et al., 2008) implemented by the function cluster_louvain in the R package igraph (version 1.0.1, (Csardi and Nepusz, 2006)). Candidates within the same communities as the original candidate sequences were used as protein candidates.

#### DNA search

To search novel peptides that were previously not annotated, we extracted all transcript sequences of the protein candidates and aligned them with muscle to generate an HMM profile (hmmbuild) that was used to search all genomes with nhmmer (3.1b2,(Wheeler and Eddy, 2013)). Candidate regions were filtered for already annotated genes and used as input to restrict *de novo* gene prediction with Augustus (version 3.3.3, (Stanke et al., 2008)). Finally, candidates from both, protein and DNA search, were merged to generate the final set of CTNIP candidates (Supplementary file 8 (CTNIP_relaxed.align)). This “relaxed” set of candidates was further filtered for having two Cysteins with a 9-11 amino acid spacing. Few candidates were also removed by a visual inspection of the alignment, resulting in the “confident” CTNIP candidates (Supplementary file 9 (CTNIP_confident.align)). Phylogeny and clade identification was done with the “relaxed” set of candidates using muscle and FastTree (version 2.1.11 SSE3, option -lg, (Price et al., 2010)) using an age cutoff of 9. The resulting phylogenetic tree was rooted using a similar sequence from *M. polymorpha* (chr5:16052258-16053618) as outgroup with gotree (v0.4.2, (Lemoine and Gascuel, 2021)) and graphically represented using FigTree (v1.4.3, http://tree.bio.ed.ac.uk/software/figtree). The sequence logo was generated with the alignment of the “confident” CTNIP candidates and the R-package ggseqlogo (v.1, (Wagih, 2017)). Amino acids with a low occurrence (i.e., seen in less than 5 % of the peptides) were trimmed from the alignment to generate a gap-free logo.

### RK identification

Protein sequences from all species were first filtered for a minimum length of 500 amino acids and merged into a single file. The initial set of RK sequences was taken from the alignment provided by Furumizu et al. (2021), but with the outgroups removed (*P. margaritaceaum*, *S. muscicola*, and *M. endlicherianum*). The sequences are given in Supplementary file 10 (SI_data_initial_RK_candidates.fasta). Sequences were aligned with muscle to build and search an HMM profile (hmmsearch options -E 1e-10 --incE 1e-10). Candidates were matched to each other with diamond (options -e 1e-11 --id 20 --query-cover 80). The pairwise percent similarity scores above 50 % were used to construct a network and communities were defined as described as above. Likewise, only candidates within the same communities as the original candidates were kept. Candidates were further filtered for the presence of an LRR and a kinase domain with hmmsearch (options -E 1e-5) and PFAMv33 (Supplementary file 11 (RECEPTOR.align)). Phylogeny and clade identification was done with muscle and FastTree (option -lg) using an age cutoff of 5.5. The resulting phylogenetic tree was rooted using the sequences from *P. margaritaceaum* as outgroup (Furumizu et al., 2021).

### Phylogenetic tree of all species

The species tree was calculated using OrthoFinder (v2.5.4, (Emms and Kelly, 2019)) with all protein sequences of all plants species.

